# Investigating Genetic Heterogeneity in Major Depression Through Item-level Genetic Analyses of the PHQ-9

**DOI:** 10.1101/528067

**Authors:** Jackson G. Thorp, Andries T. Marees, Jue-Sheng Ong, Jiyuan An, Stuart MacGregor, Eske M. Derks

## Abstract

**Background:** Major Depressive Disorder (MDD) is a clinically heterogeneous disorder. Previous large-scale genetic studies of MDD have explored genetic risk factors of MDD case-control status or aggregated sums of depressive symptoms, ignoring possible clinical or genetic heterogeneity.

**Aim:** In this study, we present the results of symptom-level genetic analyses and compare SNP-based heritability (*h*^2^ SNP) and genetic correlations across major depression symptoms. We further investigate genetic correlations with a range of psychiatric disorders and other associated traits. Methods: We have analysed data from the UK biobank and included 148,752 subjects of white British ancestry with genotype data who completed nine items of a self-rated measure of depression: the Patient Health Questionnaire (PHQ-9). Genome-Wide Association analyses were conducted for nine symptoms and two composite measures. LD score regression analysis was used to calculate SNP-based heritability (*h*^2^ SNP) and genetic correlations (r_g_) across symptoms and to investigate genetic correlations with 25 external phenotypes. Confirmatory factor analyses were applied to test whether one, two, or three-factor models best fit the pattern of genetic correlations across the nine symptoms.

**Results:** We identified 9 novel genome-wide significant genomic loci, with no overlap in loci across depression symptoms. *h*^2^ SNP ranged from 3% (suicidal ideation) to 11% (fatigue). Genetic correlations range from 0.54 to 0.96 (all *p* < 1.39×10^−3^) with 30 of 36 correlations being significantly smaller than 1. A 3-factor model provided the best fit to the genetic correlation matrix, with factors representing “psychological”, “neurovegetative”, and “psychomotor / concentration” symptoms. The genetic correlations with external phenotypes showed large variation across the nine symptoms.

**Discussion:** Patterns of *h*^2^ SNP and genetic correlations differed across the nine symptoms of depression. Our findings suggest that the large phenotypic heterogeneity observed for MDD is recapitulated at a genetic level. Future studies should investigate how genetic heterogeneity in MDD influences the efficacy of clinical interventions.

## Introduction

Clinical depression is a markedly complex and debilitating mental disorder characterised by sad, irritable or empty mood, diminished pleasure, and cognitive and somatic impairment^1^. The heritability of major depressive disorder (MDD) is estimated to be ~37% from twin studies^2^ with common Single Nucleotide Polymorphisms (SNPs) explaining around 9% of the variation in liability^3^. MDD has substantial comorbidity with other psychiatric and substance use disorders and is related to a wide range of personality, socioeconomic, and human traits^4^. There is substantial overlap in the genetic risk factors of MDD and other psychiatric disorders^3^, including significant genetic correlations (r_g_) with schizophrenia (r_g_ = 0.34), bipolar disorder (r_g_ = 0.32), autism spectrum disorders (r_g_ = 0.44) and ADHD (r_g_ = 0.42). MDD has notably high genetic overlap with anxiety disorders (r_g_ = 0.80) and neuroticism (r_g_ = 0.70), which may reflect the overlap in diagnostic criteria between the three traits. Initial efforts to identify genetic variants associated with major depression were unsuccessful, despite successes with other psychiatric diseases and traits. While a Genome Wide Association Study (GWAS) of schizophrenia (9,394 cases), for example, detected seven genome-wide significant associations ^5^, a mega-analysis of MDD (9240 cases)^6^ and a meta-analysis of depressive symptoms (N = 34,549)^7^ found no significant associations. By 2014, 108 independent genetic loci for schizophrenia had been identified^8^, and not a single one for depression. The struggle to identify significant genetic variants was likely related to low statistical power due to the clinical heterogeneity of MDD^9^.

Depression is a polygenic disorder, influenced by the combination of small effects from many genetic variants which can only be detected in studies with large sample sizes^10^. Due to the relatively high prevalence of depression (~15% vs. <1% for schizophrenia), power is lower than for other diseases with similar numbers of cases but lower prevalence^11^. Also, depression is less heritable than other psychiatric disorders (~37% vs. ~80% for schizophrenia^12^) and therefore larger sample sizes are required to obtain similar statistical power to detect significant effects. In the last two yours, increasing sample size has proved to be effective with the number of genome-wide significant variants increasing steadily with sample size. Hyde, et al. ^13^ identified 15 genome-wide significant loci associated with self-reported depression (N = 307,354). Another 17 loci were identified across three broad depression phenotypes (N = 322,580)^14^. The largest GWAS of major depression to date (N = 480,359) identified 44 significant loci^3^.

These genetic studies ignored possible clinical heterogeneity in MDD, despite clinical presentations and symptoms of MDD being diverse. The Diagnostic and Statistical Manual of Mental Disorders 5th edition (DSM-5) defines major depression by the following symptoms: (1) depressed mood, (2) diminished interest or pleasure in activities (anhedonia), (3) decrease or increase in weight or appetite, (4) insomnia or hypersomnia, (5) psychomotor agitation or retardation, (6) fatigue or loss of energy, (7) feelings of worthlessness or excessive or inappropriate guilt, (8) diminished ability to think or concentrate, or indecisiveness, and (9) recurrent thoughts of death or recurrent suicidal ideation^15^. For a diagnosis of MDD five or more of these symptoms need to be present during a two week period, with at least one symptom being depressed mood or anhedonia. Østergaard, et al. ^16^ highlighted that there are 227 possible combinations of symptoms meeting DSM-5 criteria, indicating MDD is an extremely heterogeneous disorder. Further, individual symptoms have been found to differ substantially in their association with psychosocial impairment, influence from environmental and personality risk factors, and biological correlates^17^. GWASs of depression have typically focused on MDD case-control status or aggregated sums of depressive symptoms. By combining different symptoms into a single clinical measure, it is implicitly assumed that individual symptoms of depression are genetically similar. However, the extreme heterogeneity of depression and numerous clinical presentations of the disorder suggest that different biological mechanisms could underlie the diverse subtypes of depression. Supporting this notion, depression symptoms have been found to differ substantially in heritability (h^2^ range, 0 – 35%); with somatic and cognitive symptoms being most heritable^18^. Further, the diagnostic criteria of MDD were found to reflect three underlying genetic factors (cognitive / psychomotor symptoms, mood symptoms, and neurovegetative symptoms) rather than a single factor of genetic risk in a twin study^19^. Nagel, et al. ^20^ found substantial genetic heterogeneity in neuroticism, a personality trait with extensive phenotypic and genetic overlap with MDD^21^, by conducting genetic analyses on the individual items used to measure neuroticism.

To date, it is not known to what extent genetic risk factors overlap in individual symptoms of MDD. The aim of the present study is to examine and assess the extent of genetic heterogeneity in major depression. We conduct genetic analyses on individual symptoms of depression in 148,752 participants within the UK Biobank, as measured by the nine items of the Patient Health Questionnaire (PHQ-9)^22^, a depression measure which directly maps onto the DSM-5 criteria. In order to examine genetic heterogeneity in depression we (1) conduct symptom-level GWA analyses and then compare genetic associations and SNP-based heritability across symptoms; (2) calculate phenotypic and genetic correlations across depression symptoms and determine their underlying genetic factor structure; and (3) calculate genetic correlations between individual symptoms and a range of psychiatric disorders and human complex traits.

## Methods

### UK Biobank Cohort

UK Biobank (UKBB) is a major health data resource containing phenotypic information on a wide range of health-related measures and characteristics in over 500,000 participants from the United Kingdom general population^23^. Participants were recruited between 2006 and 2010 and provided written informed consent. A total of 157,365 participants completed the PHQ-9, as part of a UKBB mental health follow-up questionnaire administered online in 2016.

### Sample selection

First, participants were included in the present study if they were of white British ancestry, identified through self-reported ethnicity and genetic principal components. Participants who self-reported as not white British, but for whom the first two genetic principal components indicated them to be genetically similar to those of white British ancestry were also included in order to maximise sample size (these commonly were participants who reported to be of Irish ancestry). Second, Participants were excluded if they were identified with schizophrenia and / or other psychotic disorders, bipolar disorder, cyclothymic disorder, or dissociative identity disorder, based on self-reported symptoms or diagnosis, reported prescription of an antipsychotic medication, and/or ICD-10 (The International Classification of Diseases, Tenth Revision) codes from linked hospital admission records. Third, only participants who provided a response for all nine items of the PHQ-9 were included (list-wise deletion represented a less than 2% reduction in sample size). This resulted in a final sample size of 148,752 (see Supplementary Figure 13 for flow diagram of sample selection).

### PHQ-9

The PHQ-9 is a commonly used self-administered measure of depression containing nine items that map directly onto the nine DSM diagnostic criteria for major depression^22^. Each PHQ-9 item assesses the frequency of that symptom over the past two weeks, rated on a four-point ordinal scale: (0) Not at all, (1) Several days, (2) More than half the days, (3) Nearly every day. (See Supplementary Table 1 for the nine symptoms of major depression, PHQ-9 items, and DSM-5 diagnostic criteria).

**Table 1:**
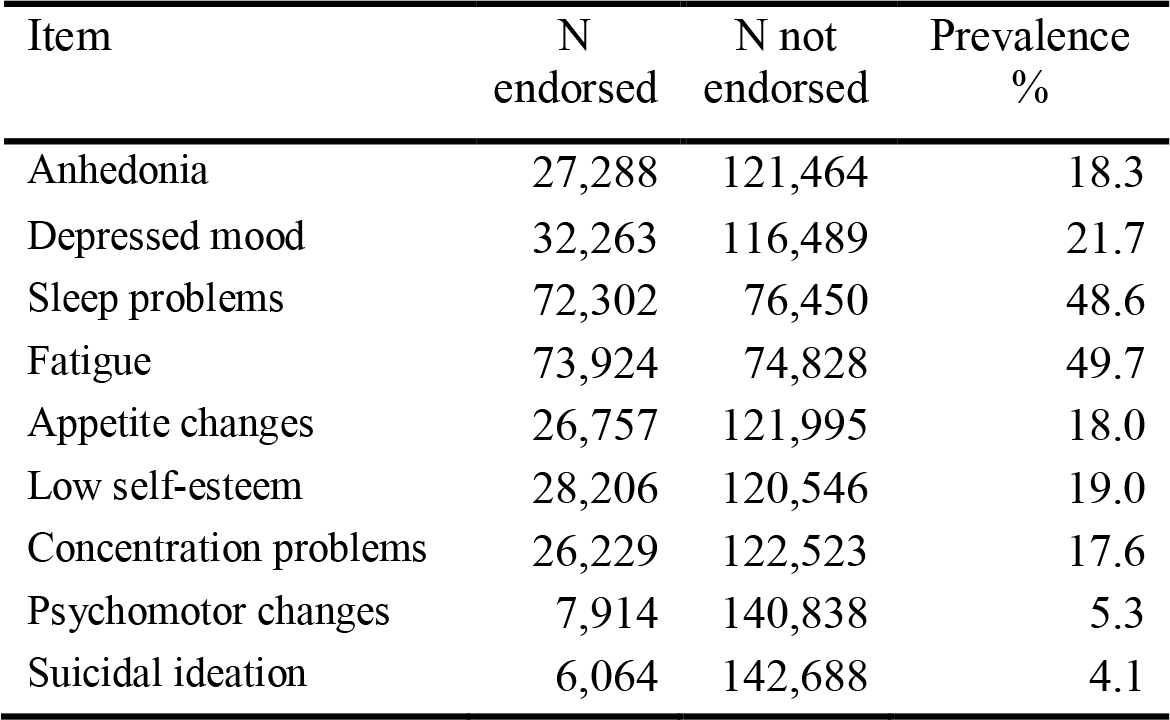
Sample sizes and prevalence of all binary PHQ-9 items.

| Item | N<br>endorsed | N not<br>endorsed | Prevalence<br>% |
| --- | --- | --- | --- |
| Anhedonia | 27,288 | 121,464 | 18.3 |
| Depressed mood | 32,263 | 116,489 | 21.7 |
| Sleep problems | 72,302 | 76,450 | 48.6 |
| Fatigue | 73,924 | 74,828 | 49.7 |
| Appetite changes | 26,757 | 121,995 | 18.0 |
| Low self-esteem | 28,206 | 120,546 | 19.0 |
| Concentration problems | 26,229 | 122,523 | 17.6 |
| Psychomotor changes | 7,914 | 140,838 | 5.3 |
| Suicidal ideation | 6,064 | 142,688 | 4.1 |

The PHQ-9 is a psychometrically valid and reliable measure of depression^24^. Test-retest reliability was high (*r* = .84, over a span of 48 hours) and internal consistency was excellent with Cronbach’s alphas (*α*) of .89 and .86 in primary care and obstetrics-gynaecology samples, respectively. The authors also reported good criterion and construct validity. The PHQ-9 was validated against professional diagnoses of MDD, resulting in 88% sensitivity and 88% specificity (at a PHQ-9 sum-score of ≥ 10); and scores correlated highly with similar constructs, such as the 20-item Short-Form General Health Survey (SF-20)^25^ mental health scale (*r* = .73). Internal consistency of the PHQ-9 in the UK Biobank sample in the current study was high (Cronbach's *α* = .83).

### Depression Item Phenotypes

Each of the nine PHQ-9 items is considered a separate phenotype in the genetic analyses. The ordinal scale of measurement of these items complicates interpretation of the SNP-based heritability estimates (amount of phenotypic variance in the item explained by SNPs). SNP-based heritability is an important concept in genetics, essential to understanding the magnitude of the genetic influence on a particular trait^26^. To enable a direct comparison across each of the PHQ-9 items, each ordinal phenotype was transformed to a binary phenotype for heritability estimation. The nine items were dichotomised such that an item was considered to be endorsed if the item score was one or greater (several days, more than half the days, or nearly every day), and not endorsed if the score was zero (not at all). A cut-off score of one was used in order to maximise the number of subjects who endorsed an item and hence statistical power, a strategy that has provided greater benefit in GWASs of depression over ensuring a seamless phenotype^3,11,14,27^. In addition to the nine ordinal items and nine binary items, a *sum-score* (sum of all ordinal item scores; ranging from 0 to 27) and *binary sum-score* (number of binary items endorsed; ranging from 0 to 9) were included as phenotypes. We will present the results from the binary items and the two sum-scores while results for ordinal items are provided in supplementary.

### Genome-Wide Association Analyses

A total of 20 GWA analyses were conducted (nine ordinal scale depression items, nine binary items, plus the sum-score and binary sum-score phenotypes) using BOLT-LMM^28^. Associations between SNPs and a phenotype are tested using a linear mixed model in order to correct for population structure and cryptic relatedness. While BOLT-LMM is based on a quantitative trait model, it can be used to analyse binary traits by treating them as continuous and applying a transformation. Ordinal items are treated as continuous. An issue when analysing binary traits in BOLT-LMM is the inflated type 1 error rates for rare SNPs when the number of cases and controls are very unbalanced^29^. In practice, all of the traits we consider here have a case proportion which is large enough (3%) for this not to be a problem^30^.

Analyses were limited to autosomal SNPs with high imputation quality score (INFO score ≥ 0.80) and a minor allele frequency of 1% or higher, resulting in 9,413,637 SNPs being tested for association. Sex and age were included as covariates. GWAS results were annotated using the FUMA GWAS platform^31^. The conventional genome-wide significance threshold of *p* < 5×10^−8^ was applied. Due to the exploratory nature of the analyses and the high correlation between the 20 phenotypes, we decided not to correct for multiple testing of the 20 phenotypes as this would lead to increased type-II error rate and reduced power.

Significant SNPs were clumped into blocks high in linkage disequilibrium (the non-random association of alleles at a specific locus; LD) using a threshold of r^2^ < 0.10 (correlation between allele frequencies of two SNPs; as calculated by PLINK). Independent significant SNPs were defined as the SNP with the lowest p-value within an LD block. Genomic risk loci (distinct, fixed positions on a chromosome) were identified by merging independent SNPs if r^2^ ≥ 0.10 and their LD blocks are physically close to each other at a distance of 1,000 kb.

### LDSC analyses

Estimates of the variance in each phenotype attributable to the additive effects of all SNPs (SNP-based heritability; *h*^2^ SNP) were calculated via single-trait LD Score Regression using GWAS summary statistics from our analyses^32^ (see Supplementary methods). In order to interpret *h*^2^ SNP for binary items estimates are converted to a normally distributed liability scale, because liability scale heritability is independent of prevalence and can be compared across different phenotypes and populations^33^. The population prevalence of PHQ-9 items was estimated from our UK Biobank sample (population prevalence = sample prevalence; see Table 1). We applied a Bonferroni corrected significance threshold for the 11 *h*^2^ SNP estimates (*p* < 4.55×10^−3^).

Cross-trait LD Score Regression^34^ was used to estimate genetic correlations (r_g_) between each of the nine binary items. We applied a Bonferroni corrected significance threshold for these 36 *r*_g_ tests (*p* < 1.39×10^−3^). Additionally, we also calculated pairwise genetic correlations between our phenotypes (9 depression items and sum-scores) and 25 other psychiatric, substance use, socioeconomic and human traits with publicly available GWAS summary statistics (see Supplementary Table 2). Multiple testing was corrected for by adjusting p values based on false discovery rate (FDR) across all tests.

**Table 2:** GWAS Results for binary PHQ-9 items and sum-score phenotypes.

| Gen. locus | SNP id | Chr | BP | A1 | A2 | Freq A1 | Phenotype | $\beta$ | SE | p-value | nSNPs | nearest gene | eQTL genes |
| --- | --- | --- | --- | --- | --- | --- | --- | --- | --- | --- | --- | --- | --- |
| 1 | rs2279681 | 1 | 201861016 | C | G | 0.657 | Sum-score | 0.093 | 0.014 | 5.60E-11 | 25 | *SHISA4* | SHISA4, LMOD1 |
|  |  |  |  |  |  |  | Binary sum-score | 0.023 | 0.004 | 8.60E-09 | 6 | *SHISA4* |  |
| 2 | rs62158169 | 2 | 114081827 | C | T | 0.784 | Sleep problems | 0.015 | 0.002 | 1.40E-10 | 22 | *PAX8* | FOXD4L1, CBWD2 |
| 3 | rs137997194 | 3 | 48824937 | A | G | 0.955 | Depressed mood | -0.023 | 0.004 | 7.70E-10 | 15 | *PRKAR2A* | AMT, KLHDC8B |
|  | rs143756010 | 3 | 49312248 | C | T | 0.956 | Binary sum-score | -0.053 | 0.009 | 7.40E-09 | 15 | *C3orf62* | AMT, NICN1, RNF123 |
| 4 | rs12492113 | 3 | 50521402 | G | A | 0.875 | Binary sum-score | -0.037 | 0.006 | 1.20E-10 | 114 | *CACNA2D2* | C3orf18, CACNA2D2, CYB561D2, DOCK3, HEMK1, HYAL1, LSMEM2, RBM6, MANF, MAPKAPK3 |
|  |  |  |  |  |  |  | Sum-score | -0.124 | 0.020 | 1.30E-09 | 76 | *CACNA2D2* |  |
|  |  |  |  |  |  |  | Appetite changes | -0.013 | 0.002 | 8.80E-09 | 28 | *CACNA2D2* |  |
| 5 | rs13127129 | 4 | 30501860 | A | G | 0.499 | Sleep problems | 0.010 | 0.002 | 4.90E-08 | 1 | *PCDH7* | - |
| 6 | rs7073667 | 10 | 107809043 | T | C | 0.452 | Sleep problems | 0.010 | 0.002 | 4.30E-08 | 12 | *RP11-298H24.1* | - |
| 7 | rs140920627 | 11 | 41577268 | C | A | 0.988 | Anhedonia | -0.037 | 0.007 | 3.70E-08 | 1 | *RP11-124G5.3* | - |
| 8 | rs840161 | 12 | 57323523 | A | G | 0.366 | Binary sum-score | -0.021 | 0.004 | 3.90E-08 | 1 | *SDR9C7* | - |
| 9 | rs2335859 | 19 | 7871837 | T | G | 0.602 | Anhedonia | -0.009 | 0.001 | 9.80E-09 | 10 | *EXOSC3P2* | - |

Note: Table displays SNPs significant at $p < 5 \times 10^{-8}$ and independent at $r^2 < 0.10$ . Genomic risk loci (Gen. locus) are defined by $r^2 < 0.10$ , window size 1,000kb. Chromosome (Chr), location in base pairs (BP) on Hg19, effect allele (A1), allele 2 (A2), frequency of effect allele (Freq A1), effect size beta ( $\beta$ ), standard error of beta (SE), p-value, and number of SNPs clumped under lead SNP (nSNPs) are shown. Proximity (nearest gene) and eQTL (eQTL genes) mapping results are given. eQTL mapping limited to significant ( $FDR < .05$ ) cis-eQTLs from GTEx v7<sup>53</sup> and the CommonMind Consortium (CMC)<sup>54</sup>. SNP rs137997194 and rs143756010 are tagging the same signal, but was the SNP with lowest p-value in “depressed mood” and “binary sum-score”, respectively.

### Hierarchical Cluster Analysis

A hierarchical cluster analysis was conducted to examine the underlying genetic structure between depression items. Implemented in the hclust function in R^35^, items are grouped into similar clusters based on a measure of dissimilarity between each pair of items and the results are presented in a cluster dendrogram. Dissimilarity was defined as one minus the genetic correlation (1 − *r*_g_).

### Confirmatory Factor Analyses

Confirmatory factor analyses (CFA) were conducted based on genetic covariances between items, in order to quantitatively assess the genetic factor structure of the PHQ-9 identified in the cluster analysis. The fit of a one-factor baseline model and two and three-factor models identified in the cluster analysis were compared.

*χ*^2^ likelihood ratio tests are very sensitive to large samples and often produce spurious positive results^36^. Given the very large sample size in the present study, model fit was evaluated with a range of alternative fit indices. These indices (and their commonly used thresholds for acceptable model fit) include: NFI (≥ .95), AGFI (≥ .95), RMSEA (≤ .06), and SRMR (≤ .06)^37^. Models were compared using AIC and BIC indices, which take into account both model fit and complexity. The most parsimonious model is the model with the lowest AIC and BIC values.

## Results

### Descriptive Statistics

The final sample (N = 148,752) was 56% female, ranging in age from 38 to 72 years old (*M* = 55.93, *SD* = 7.73). The distribution of responses to all PHQ-9 items (on the ordinal scale) are displayed in Supplementary Table 3. The distribution of item scores varied considerably across items; sleep problems and fatigue had the highest endorsement rates while suicidal ideation and psychomotor changes had the lowest rates. Sum-scores ranged from 0 to 27, with a mean of 2.71 (*SD* = 3.61). Endorsement rates of binary depression items are shown in Table 1. The number of symptoms endorsed ranged from zero to nine, with a mean of 2.02 (*SD* = 2.20).

### GWA Analyses

Genome-wide association analyses of the 9 binary depression items plus sum-score phenotypes identified a total of 326 genome-wide significant SNPs (*p* < 5×10^−8^), tagged by 13 independent SNPs. Two lead SNPs were significant in more than one phenotype, such that across all phenotypes there are 11 unique, independent genome-wide significant SNPs. These SNPs mapped onto nine genomic risk loci (see Table 2 for results, Supplementary Figures 1-10 for QQ plots and Manhattan plots of all phenotypes; and Supplementary Table 4 for the ordinal item GWAS results).

### Heritability Estimates

Estimates of the proportion of phenotypic variance in each item attributable to the additive effects of all SNPs (SNP-based heritability; *h*^2^ SNP) varied considerably across the nine items (see Figure 1 and Supplementary Table 5). All estimates were significant after Bonferroni correction (*p* < 4.55×10^−3^). The amount of variance explained by common SNPs ranged from 3% of variance in suicidal ideation up to 11% of the variance in fatigue (mean *h*^2^ SNP across the nine depression items was 7%). *h*^2^ SNP estimates for the sum-score and no. symptoms phenotypes were 6% and 7%, respectively.

**Figure 1.**
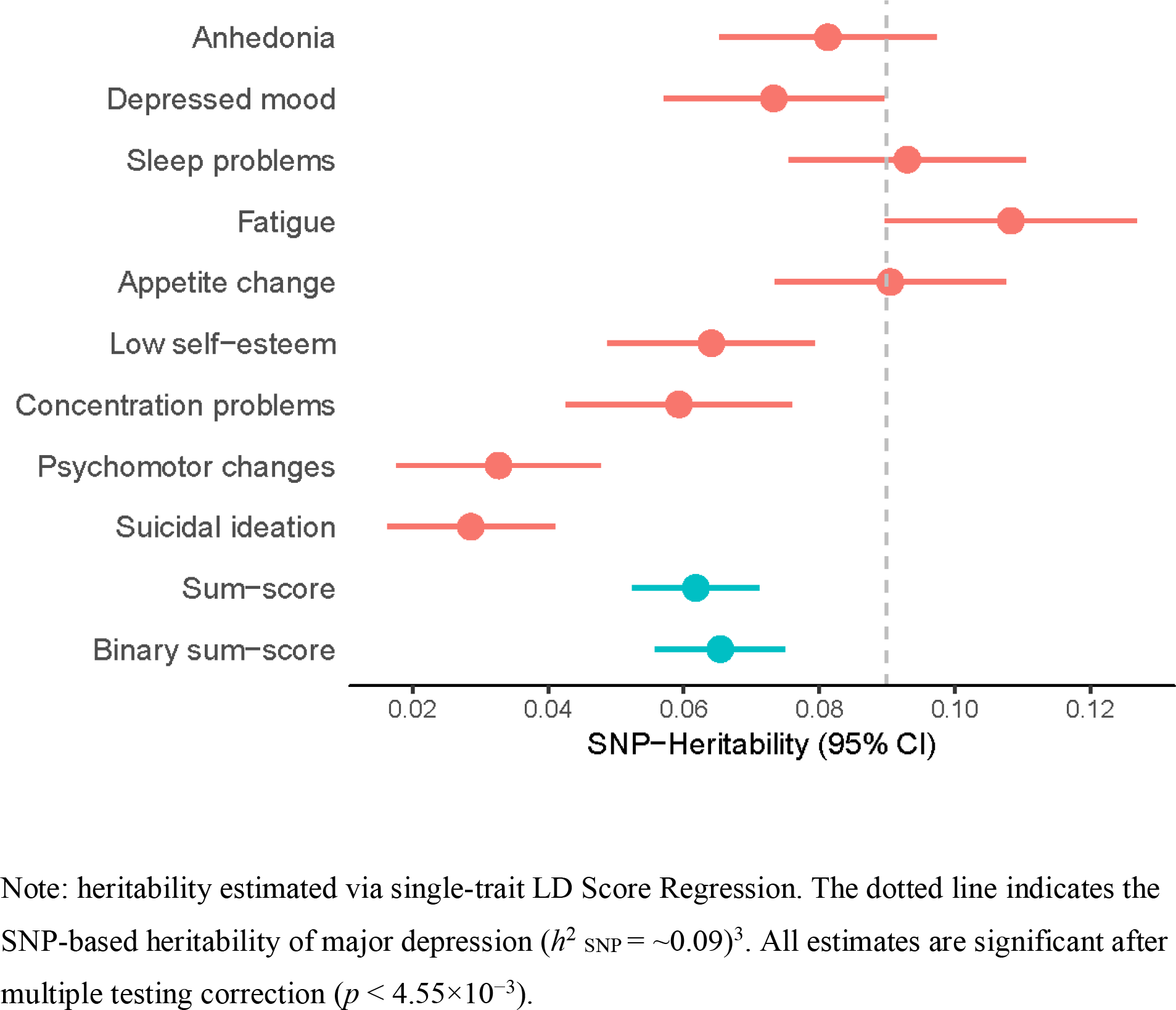
SNP-based heritability estimates and 95% confidence intervals (95% CI) for the nine depression items and sum-score phenotypes.

### Inter-item Phenotypic and Genetic Correlations

Spearman correlations between all pairs of PHQ-9 depression items showed that all items were positively correlated with each other phenotypically and remained significant after Bonferroni correction for 36 tests (*p* < 1.39×10^−3^). Coefficients ranged from .19 to .69, with the strongest association between anhedonia and depressed mood, the two core symptoms of MDD (see Figure 2).

**Figure 2.**
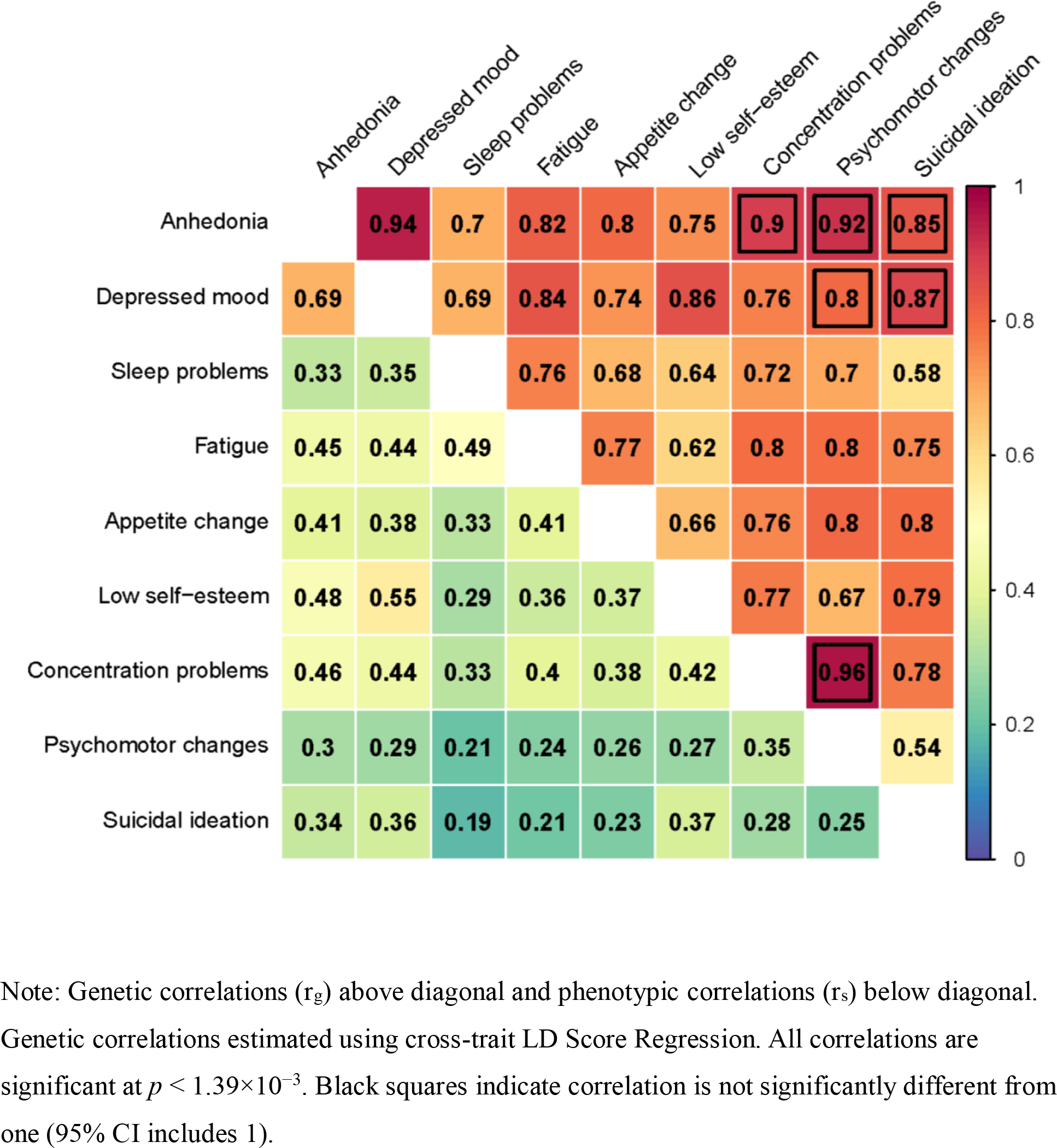
Inter-item genetic and phenotypic correlations.

Summary statistics from the GWASs of the nine binary items were used to calculate genetic correlations (r_g_) between items. All correlations were significant after correcting for multiple testing (*p* < 1.39×10^−3^) and were in the same direction (see Figure 2). Estimated r_g_’s ranged from .54 (suicidal ideation / psychomotor changes; *s.e* = .15) to .96 (psychomotor changes / concentration problems; *s.e* = .11), with a mean r_g_ of .77. Thirty out of the 36 genetic correlations were significantly less than one (95% CI did not include one), indicating substantial genetic heterogeneity across the PHQ-9 items (partly unique genetic risk factors contribute to the majority of pairs of depressive symptoms; see Figure 2 and Supplementary Table 6). Some of the genetic correlations that were not significantly different from 1 were relatively low, but have large standard errors which explains their overlap with 1.

A very similar pattern of genetic correlations emerged for the ordinal items (r_g_ range: .55 to .96), such that the Pearson correlation between the set of binary item r_g_’s and ordinal item r_g_’s was high, *r* = .90, *p* < .001 (see Supplementary Figure 11).

The Pearson correlation between the genetic correlations and phenotypic correlations was moderate, *r* = .48, *p* = .003, suggesting phenotypic correlations do not map one to one with genetic correlations (see Supplementary Figure 12).

### Genetic Clustering Analysis

A hierarchical clustering analysis based on genetic covariance between the nine depression items revealed two main genetic clusters: the first cluster including anhedonia, depressed mood, suicidal ideation, and low self-esteem (psychological symptoms); and the second cluster including psychomotor changes, concentration problems, fatigue, appetite change, and sleep problems (somatic symptoms; see Figure 3). Further exploration of the cluster dendrogram suggests the somatic symptoms cluster could again be split into two clusters: “neurovegetative” symptoms (fatigue, appetite change, and sleep problems); and “psychomotor / concentration” symptoms.

**Figure 3.**
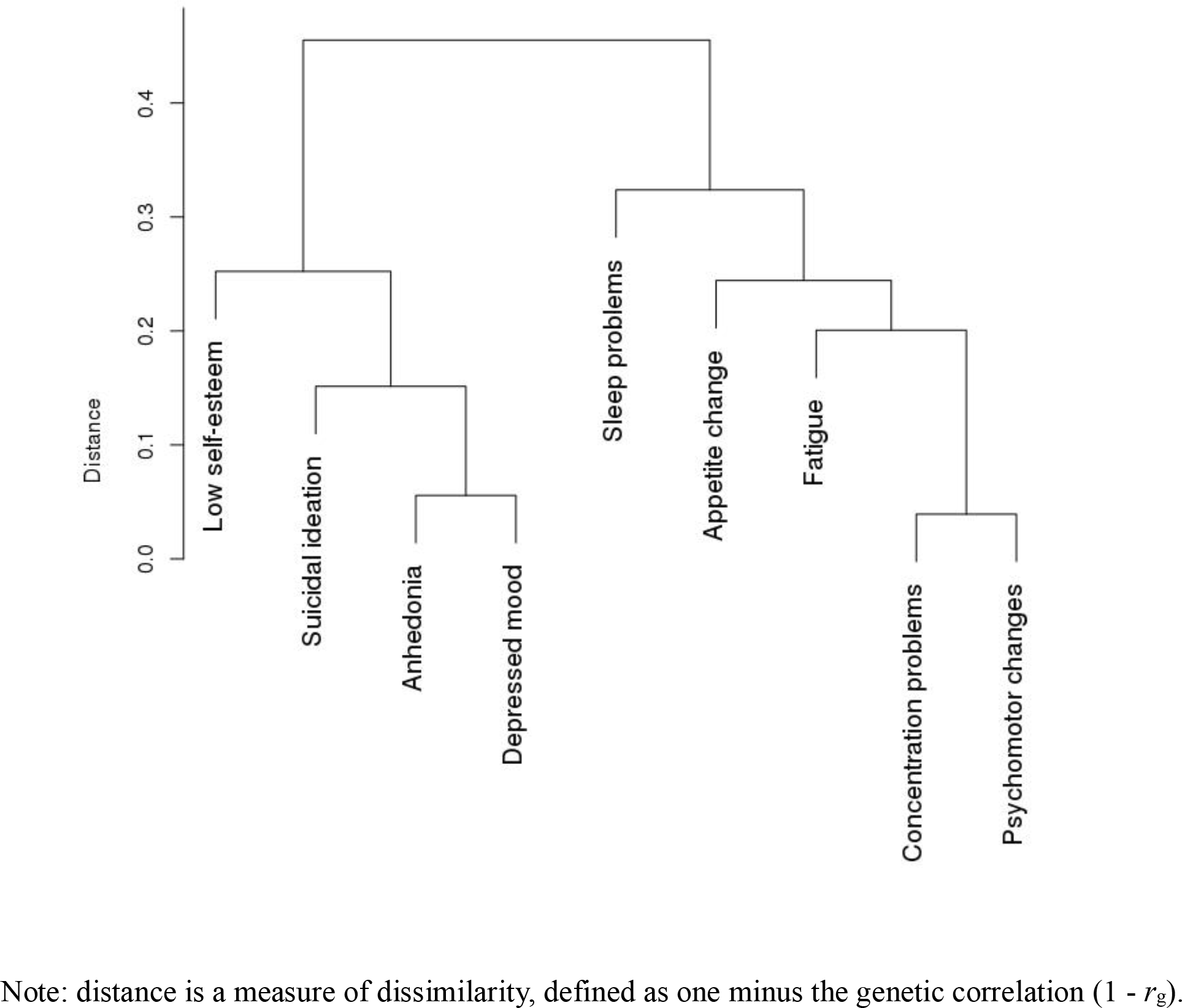
Cluster dendrogram of inter-item genetic correlations.

### Confirmatory Factor Analyses

CFA of the genetic factor structure found that all three models provided good fit to the data (see Supplementary Table 7). Comparison of models based on AIC and BIC values found that the three-factor model was the most parsimonious model compared to the one-factor model and the two-factor model (substantially lower AIC and BIC values). These results suggest the PHQ-9 is reflected genetically by three factors, comprising “psychological”, “neurovegetative”, and “psychomotor / concentration” symptoms.

### Genetic Correlations with External Traits

Genetic correlations of the nine depression items, sum-score and binary sum-score with 25 other psychiatric, substance use, socioeconomic and human traits are displayed in Figure 4. Correlations significant after correcting for false discovery rate (FDR) are indicated by non-white squares (see Supplementary Table 8).

**Figure 4.**
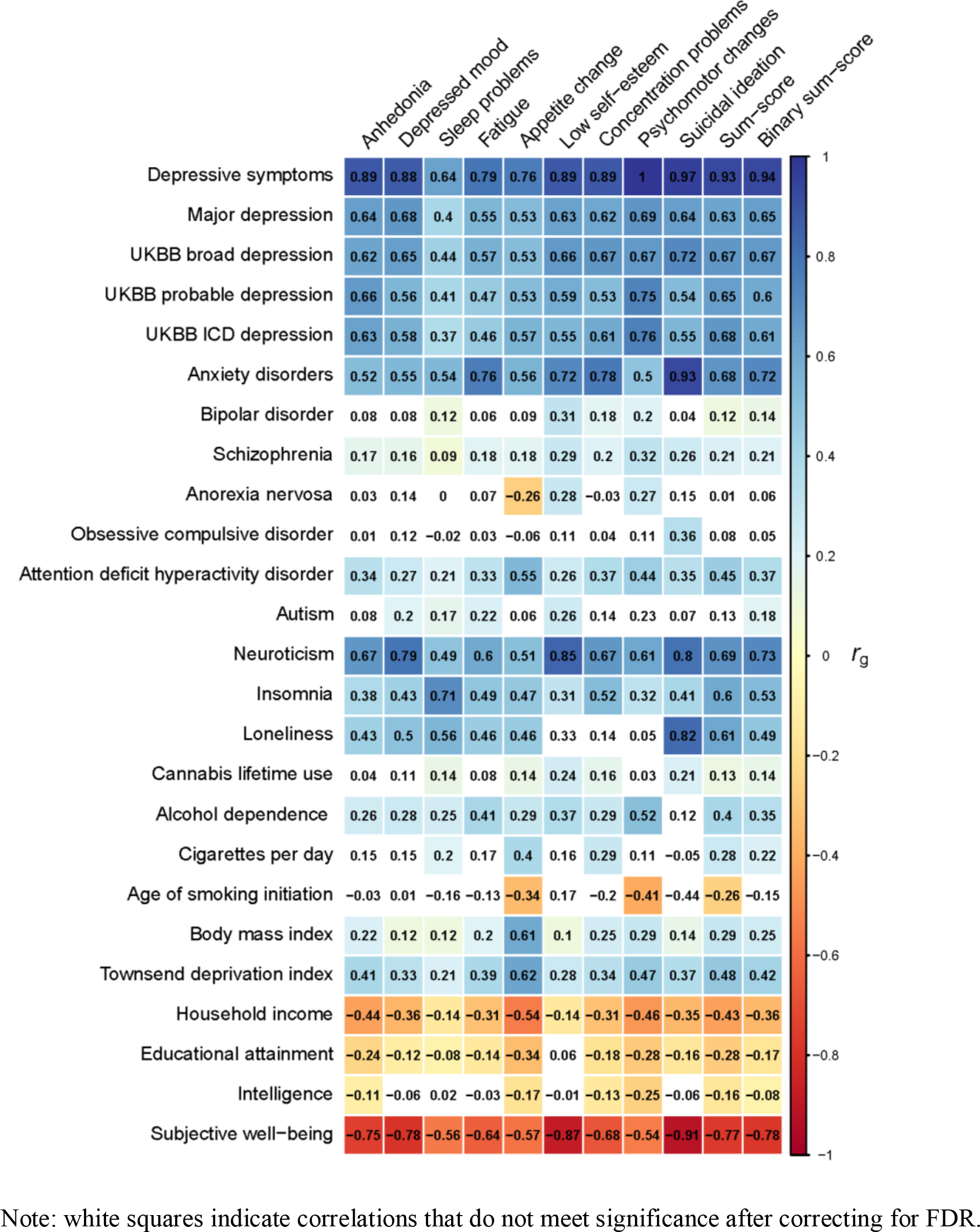
Genetic correlations between PHQ-9 items and a range of other complex traits (psychiatric, substance use, and socioeconomic phenotypes) based on publicly available summary statistics.

Individual depression items correlated as expected with closely related traits, supporting the validity of the individual symptom phenotypes in the present study. For example, appetite change had a substantially stronger positive genetic correlation with body mass index (*r*_g_ = .61) than the other eight depression symptoms (*r*_g_’s range between .10 to .29); and sleep problems had a strong, positive correlation with insomnia (*r*_g_ = .71). All symptoms were negatively correlated with subjective well-being (*r*_g_ range = −.54 to −.91), with suicidal ideation having the strongest association. Furthermore, all items positively correlated (and showed a similar pattern) with the other MDD and overall depression phenotypes.

Genetic overlap with other psychiatric disorders and traits differed substantially across depression symptoms, such as with anxiety disorders (*r*_g_ range = .50 to .93), neuroticism (*r*_g_ range = .49 to .85), schizophrenia (*r*_g_ range = .09 to .32), and insomnia (*r*_g_ range = .31 to .71). Furthermore, bipolar disorder was significantly correlated with 4 out of 9 depression items only (sleep problems, low self-esteem, concentration problems, and psychomotor changes). Anorexia nervosa overlapped with just three items, with genetic correlations even being in different directions (low self-esteem *r*_g_ = .28, psychomotor changes *r*_g_ = .27, and appetite change *r*_g_ = −.26).

## Discussion

In the present study, we investigated genetic heterogeneity in major depression by conducting genetic analyses on individual symptoms of MDD in 148,752 participants from the UK Biobank. We identified nine genomic risk loci across the nine MDD symptoms and sum-score phenotypes, all have not been associated with major depression in previous GWASs^3,7,13,14,27,38–43^. Our results revealed substantial genetic heterogeneity in depression symptoms with no overlap in significant loci across PHQ items. Though we acknowledge that the lack of overlap may be due to low statistical power to detect all true associations, we highlight some notable examples where a specific symptom of depression is linked to a gene that was previously found to be associated with a strongly related phenotype. For the item “sleep problems”, we found SNPs that implicate *PAX8* (based on proximity), a transcription factor related to thyroid follicular cell development and expression of thyroid-specific genes, replicating previous studies linking this gene to sleep duration^44–46^. In addition, SNPs associated with “depressed mood” influenced the expression of *KLHDC8B* (protein coding gene involved in cytokinesis). This gene has been previously linked to depressed affect, a sub-cluster of neuroticism that is strongly related to depression^47^. Neither of these genes were implicated in the largest GWASs of overall depression^3,14^, illustrating the importance of exploring genetic associations for specific symptoms of depression.

SNP-based heritability analyses revealed that individual depression symptoms were differentially heritable (*h*^2^ SNP ranging from 3 to 11%), suggesting that depression symptoms differ in their relative proportions of common SNP contributions. Notably, items within the “neurovegetative” symptom cluster were most highly heritable, consistent with a previous report that found somatic symptoms (such as sleep problems and appetite changes) to have a stronger heritable basis ^18^.

Genetic correlations between depression symptoms ranged from moderate (*r*_g_ < .60) to high (*r*_g_ > .90), suggesting that while some symptoms have high genetic overlap, a substantial amount of genetic variation is not shared between symptoms. This indicates extensive genetic heterogeneity in major depression, in line with the finding that depression represents multiple dimensions of genetic risk^19^ and previous associations between individual symptoms and specific polymorphisms^48^.

The underlying genetic structure between symptoms was best explained by three genetic clusters. This suggests there are risk factors specific to clusters which could indicate underlying biology specific to either “neurovegetative”, “psychological” or “psychomotor / concentration” symptoms of depression. This is consistent with symptoms differing in their biological correlates, with neurovegetative symptoms such as weight gain, increased appetite, and sleep problems being associated with higher levels of inflammation markers^49,50^. These clusters were not in full agreement with the three genetic factors found by Kendler, et al. ^19^ based on an analysis of twin data. As an example, in the Kendler study, suicidal ideation loaded onto the same factor as psychomotor changes and concentration problems, while we find that suicidal ideation clusters with symptoms of depressed mood, anhedonia, and low self-esteem that together form a “psychological symptoms” factor. However results are not easily comparable given that they derived factors from a twin study (and therefore captured rare genetic variants as well as common SNPs), used a subset of eight symptoms (appetite changes did not load onto any factor), and symptom phenotypes came from structured clinical interview rather than a self-report measure such as the PHQ-9.

Results from genetic correlations between items and a range of external traits lead us to note three general observations. First, genetic correlations with external traits differed substantially between symptoms providing evidence for genetic heterogeneity in major depression. In agreement with previous findings for major depression^3,14^, all symptoms overlapped with anxiety, schizophrenia, ADHD, insomnia, neuroticism, and subjective well-being, however the proportion of overlap varied considerably across symptoms. For example, anxiety disorders had a substantially higher genetic correlation with “suicidal ideation” than the other items. This supports a strong phenotypic association, with over 70% of people with a history of suicide attempt having an anxiety disorder, compared to ~33% in the general population^51^. Second, some traits (such as bipolar disorder, cannabis lifetime use, cigarettes per day, and intelligence) were genetically correlated with a subset of items only. Bipolar disorder for example, was genetically correlated with only four items (low self-esteem, concentration, psychomotor changes, and sleep problems), suggesting that the moderate genetic overlap between bipolar and major depression^52^ is predominately driven by these selected symptoms. This highlights how insight into the genetic architecture between traits can be gained from conducting symptom-level analyses. Third, we found traits that were genetically correlated with individual items, but not with the sum-score phenotypes. Anorexia nervosa did not overlap with aggregate measures of depression symptoms as operationalized in the sum-score phenotypes, in agreement with Howard, et al. ^14^ who similarly found no genetic overlap between anorexia and their three overall depression phenotypes. Yet, anorexia nervosa was genetically correlated with appetite change, low self-esteem, and psychomotor changes. Interestingly, the overlap with appetite change was in the opposite direction compared to the other two items. This finding emphasises the importance of analysing individual symptoms of a disorder, as important information is ignored by relying on sum-scores or overall phenotypes.

### Limitations

The findings and conclusions of this study should be interpreted in view of some key limitations. First, despite having the largest sample available to date, the current study is still underpowered to detect significant SNPs. Given the relatively high prevalence and phenotypic heterogeneity of depression, much larger sample sizes are needed compared to other psychiatric disorders^11^. To not reduce power further we did not correct for multiple testing (of 11 GWA analyses) and hence our GWAS results require independent replication. Second, depression items were analysed in isolation, regardless of the overall MDD status of the participant. For example, a participant could strongly endorse the symptom fatigue, yet have no other signs of depression and hence the endorsement of fatigue is unrelated to major depression. Nevertheless, it is possible that fatigue, regardless of the context it occurs in, possesses the same underlying genetic basis. Third, we used a PHQ-9 cut-off score of 1 to dichotomise items in order to maximise the number of cases and improve statistical power. A PHQ-9 item score of one does not meet the diagnostic criteria for endorsement, hence the phenotypes may represent a predisposition to rather than full endorsement of the particular symptom. Fourth, our results may be affected by ascertainment bias due to healthy volunteerism within the UKBB. As such our sample could represent a truncated version of the population’s genetic distribution for symptoms (people on far end of liability scale may be less likely to participate), hence resulting in reduced number of cases for some symptoms or reduced variation between cases and controls.

### Implications

The recent success in the discovery of genetic variants associated with depression has been driven by ever increasing sample sizes, an approach that has been favoured over reducing phenotypic heterogeneity. Consequently, GWASs have been conducted on a diverse range of depression-related phenotypes that often include a small subset of symptoms, generally with the view that the increase in sample size can overcome the lack of clinical precision. While this has indeed been proven to be effective at increasing the number of significant variants identified, our finding of symptom-level genetic heterogeneity raises questions about this approach. Using broad diagnostic phenotypes ignores the unique genetic factors associated with specific symptoms of depression that would likely provide useful information to further unravel the genetic architecture of MDD. Further, our finding of genetic heterogeneity across MDD symptoms implicates that patients with MDD show variation in disease pathogenesis. This variation may be linked to response to clinical interventions, such that patients presenting with specific symptom patterns (e.g., characterized primarily by neurovegetative symptoms) may be expected to respond differently.

### Conclusion

Our results provide convincing evidence that major depression is a genetically heterogeneous disorder, and highlight the utility of analysing the genetics of individual items or symptoms of a psychiatric disorder. Insights into the genetic aetiology and underlying biology of MDD will be maximised by combining large-scale genetic studies of broad clinical definitions with follow-up studies of more refined phenotypic measures of specific diagnostic subtypes.

## Acknowledgments

This work was conducted using the UK Biobank Resource (application number 25331). The UK Biobank was established by the Wellcome Trust medical charity, Medical Research Council (UK), Department of Health (UK), Scottish Government, and Northwest Regional Development Agency. It also had funding from the Welsh Assembly Government, British Heart Foundation, and Diabetes UK.

S.M. is supported by a National Health and Medical Research Council (NHMRC) Fellowship. A.T.M. is supported by the Foundation Volksbond Rotterdam.

