## Supplementary Figures for "Investigating Genetic Heterogeneity in Major Depression Through Item-level Genetic Analyses of the PHQ-9"

**Supplementary Figures 1 – 13**


*Figure 1.* Anhedonia Manhattan and QQ plots

*Figure 2.* Depressed mood Manhattan and QQ plots

*Figure 3.* Sleep problems Manhattan and QQ plots

*Figure 4.* Fatigue Manhattan and QQ plots

*Figure 5.* Appetite changes Manhattan and QQ plots

*Figure 6.* Low self-esteem Manhattan and QQ plots

*Figure 7.* Concentration problems Manhattan and QQ plots

*Figure 8.* Psychomotor changes Manhattan and QQ plots

*Figure 9.* Suicidal ideation Manhattan and QQ plots

*Figure 10.* Sum-score Manhattan and QQ plots

*Figure 11.* Comparison of inter-item genetic correlations between binary items and ordinal items

*Figure 12.* Scatterplots of genetic correlations vs. phenotypic correlations for both ordinal and binary items

*Figure 13.* Flow diagram of exclusions / inclusions leading to final sample size

*Supplementary Figure 1.* **Anhedonia** Manhattan and QQ plots (binary phenotype above and ordindal below)

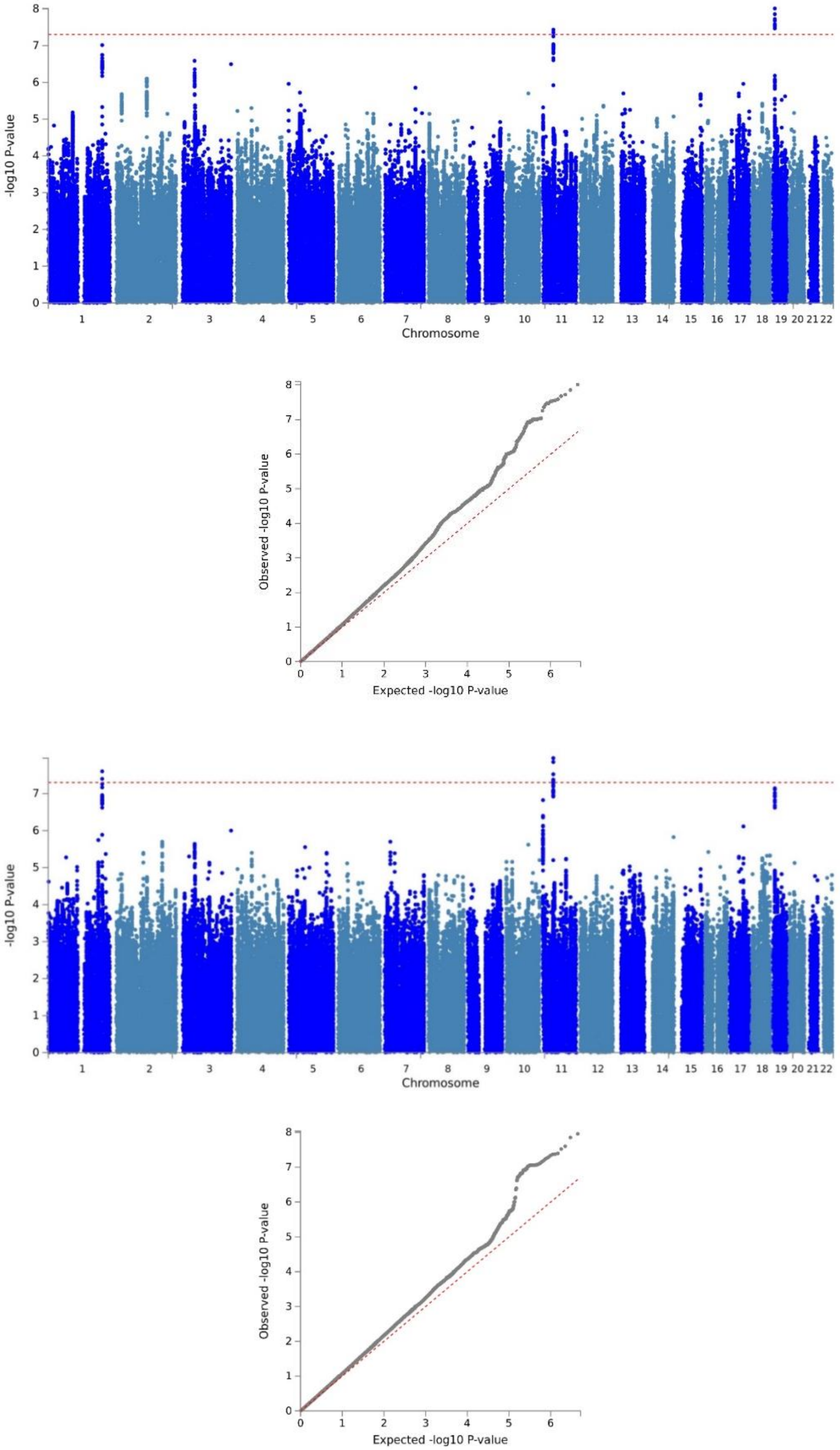

*Supplementary Figure 2.* **Depressed mood** Manhattan and QQ plots (binary phenotype above and ordindal below)

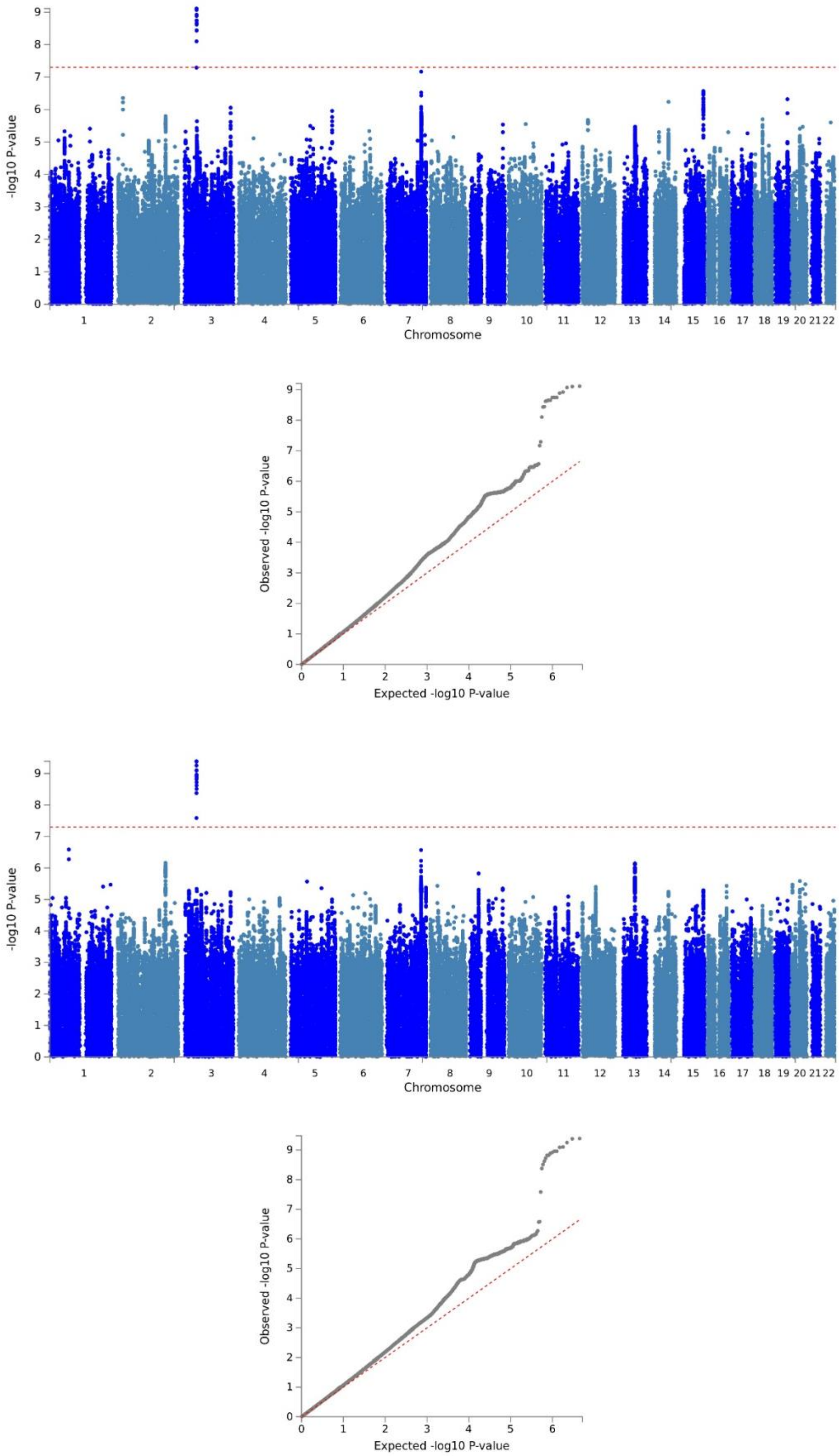

*Supplementary Figure 3.* **Sleep problems** Manhattan and QQ plots (binary phenotype above and ordindal below)

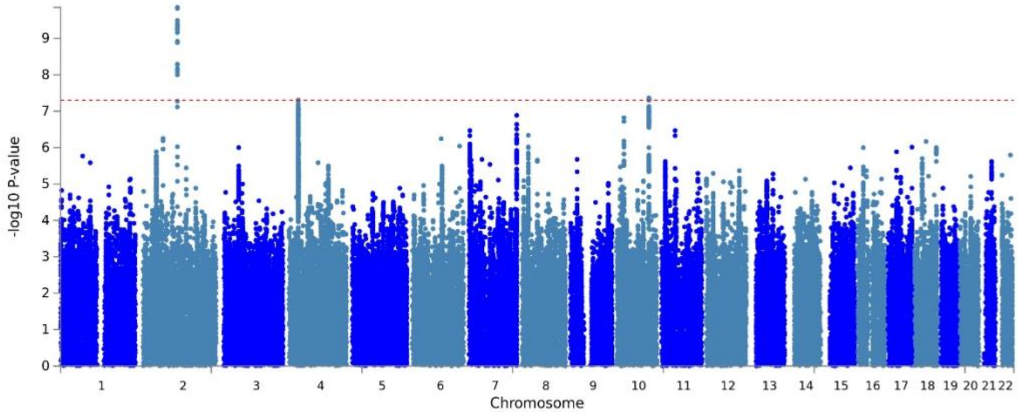


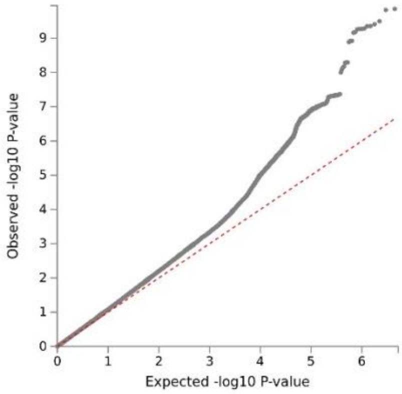


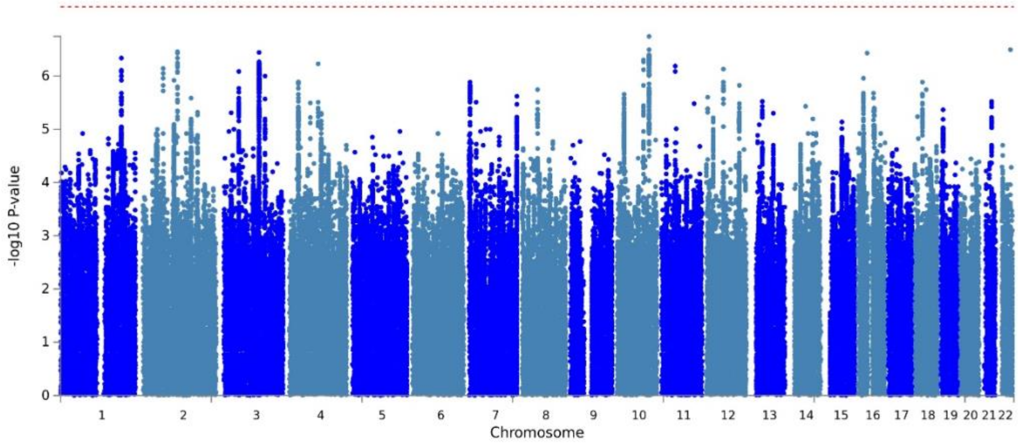


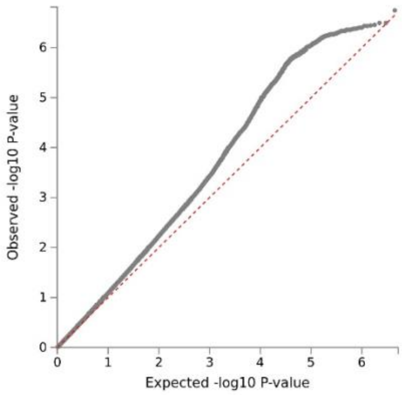

*Supplementary Figure 4.* **Fatigue** Manhattan and QQ plots (binary phenotype above and ordindal below)

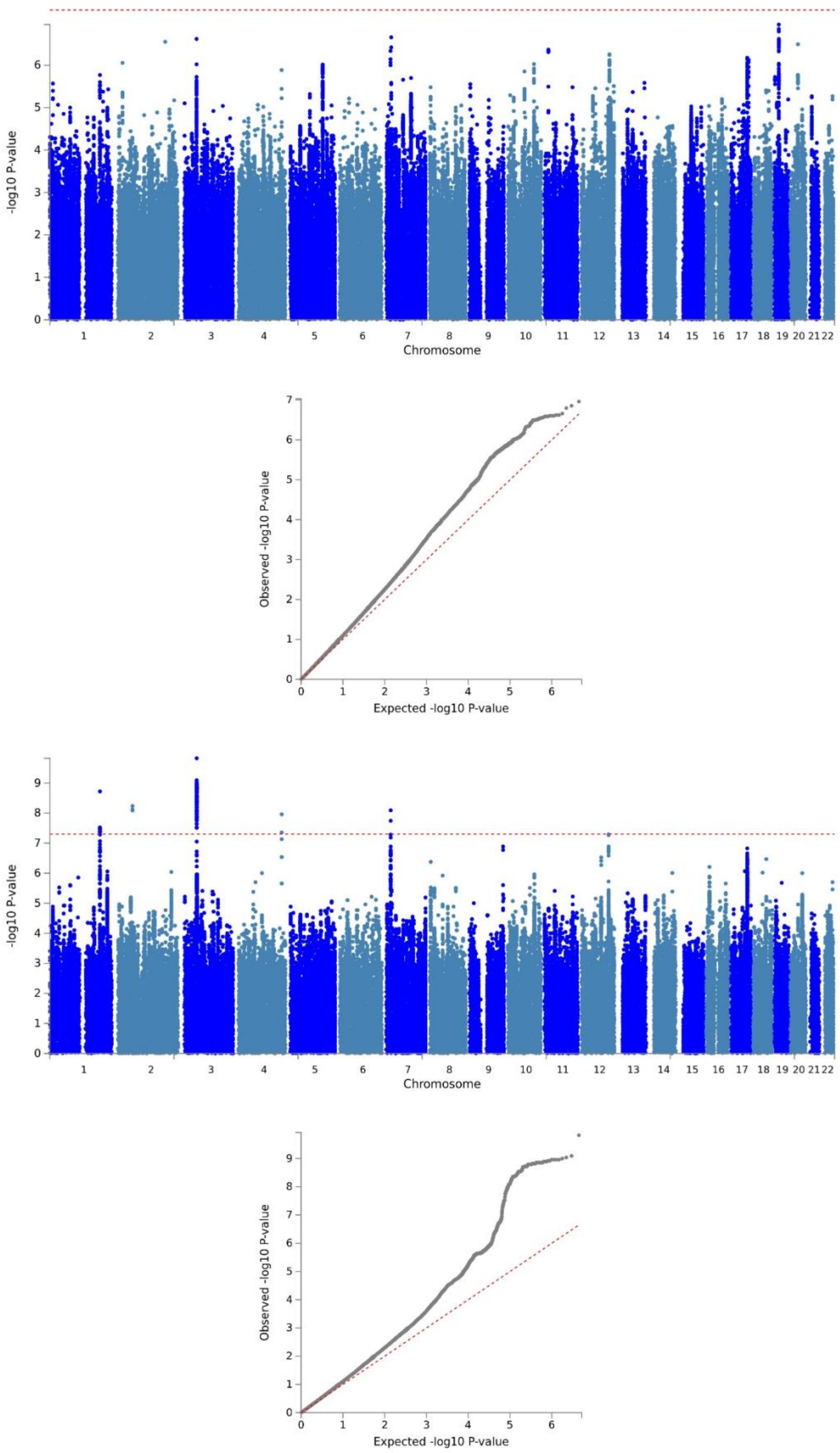

*Supplementary Figure 5.* **Appetite changes** Manhattan and QQ plots (binary phenotype above and ordindal below)

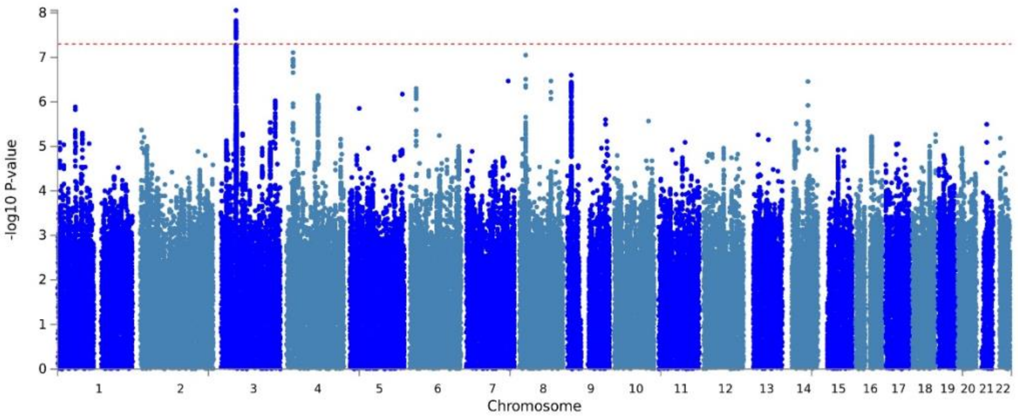


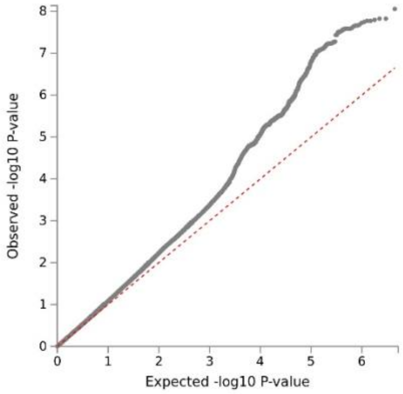


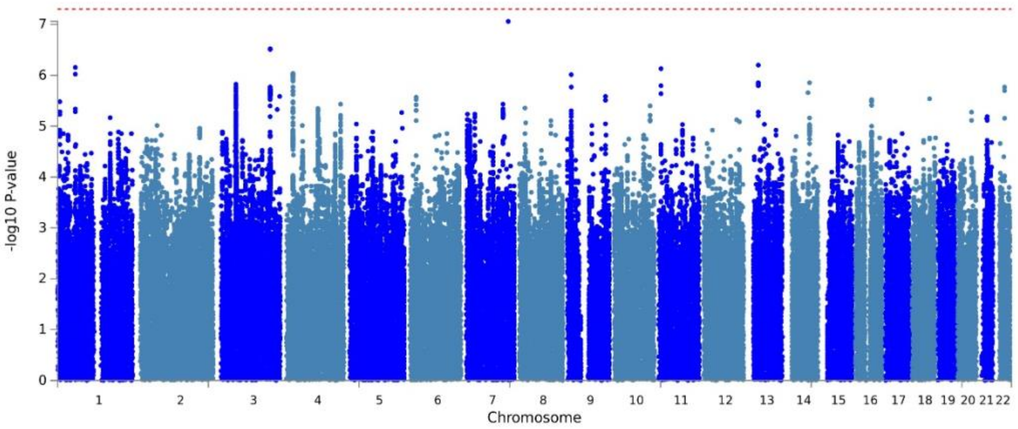


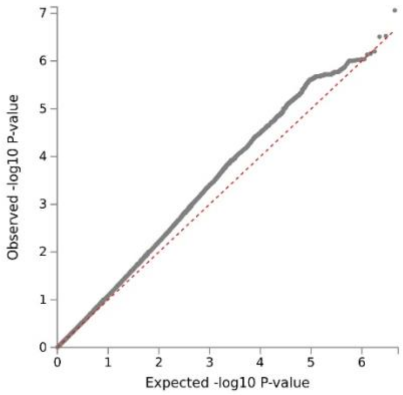

*Supplementary Figure 6.* **Low self-esteem** Manhattan and QQ plots (binary phenotype above and ordindal below)

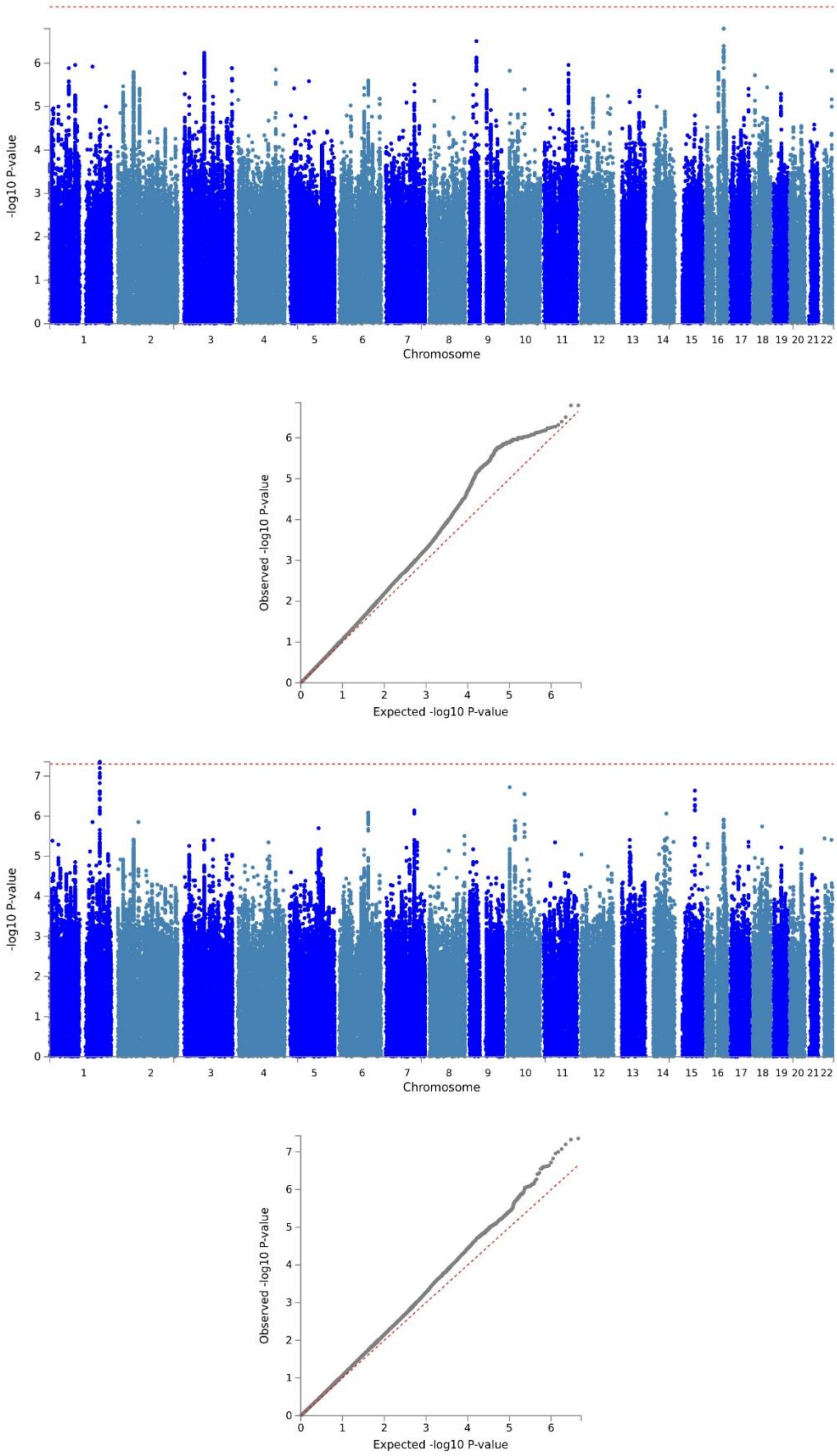

*Supplementary Figure 7.* **Concentration problems** Manhattan and QQ plots (binary phenotype above and ordindal below)

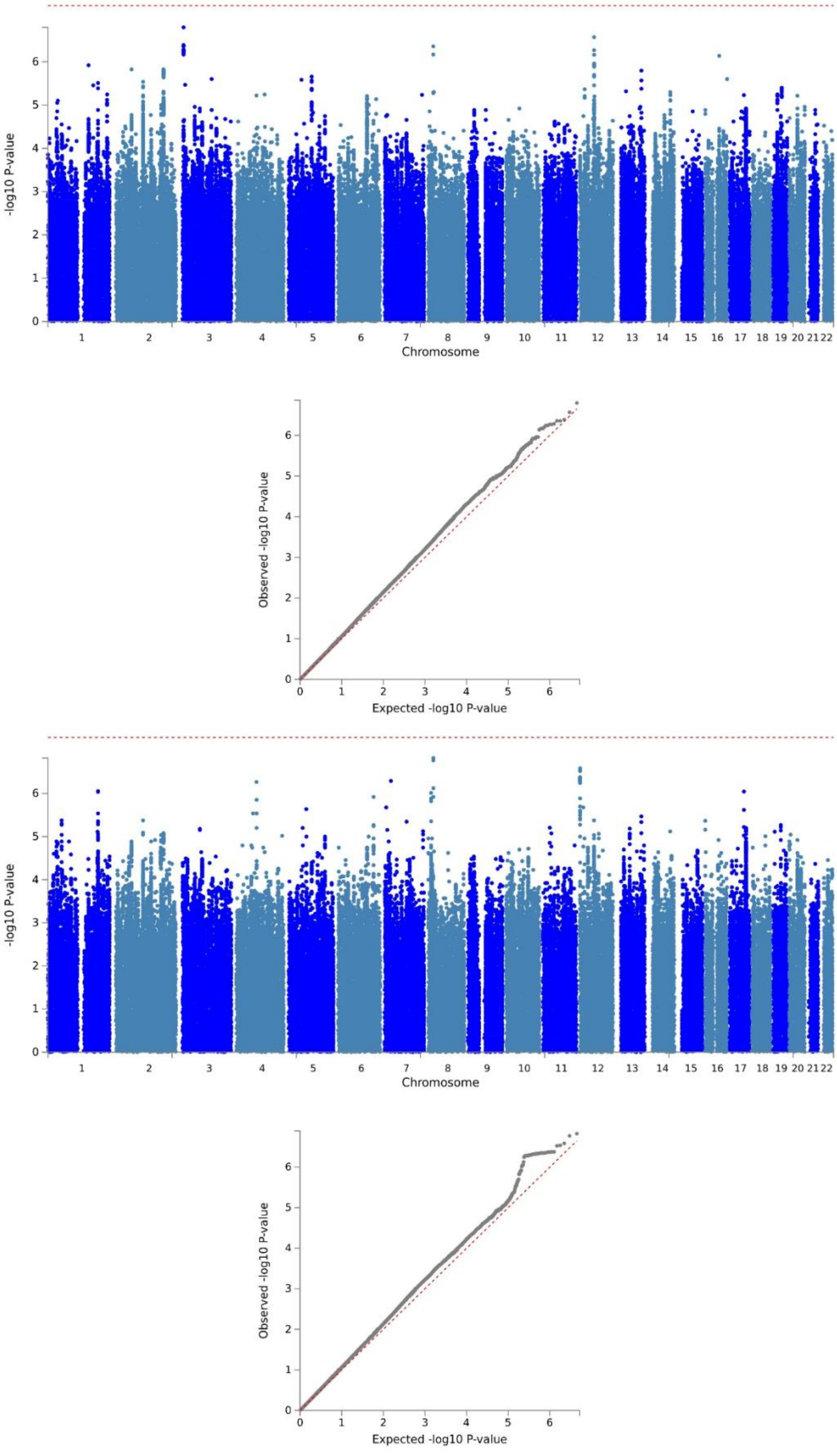

*Supplementary Figure 8.* **Psychomotor changes** Manhattan and QQ plots (binary phenotype above and ordindal below)

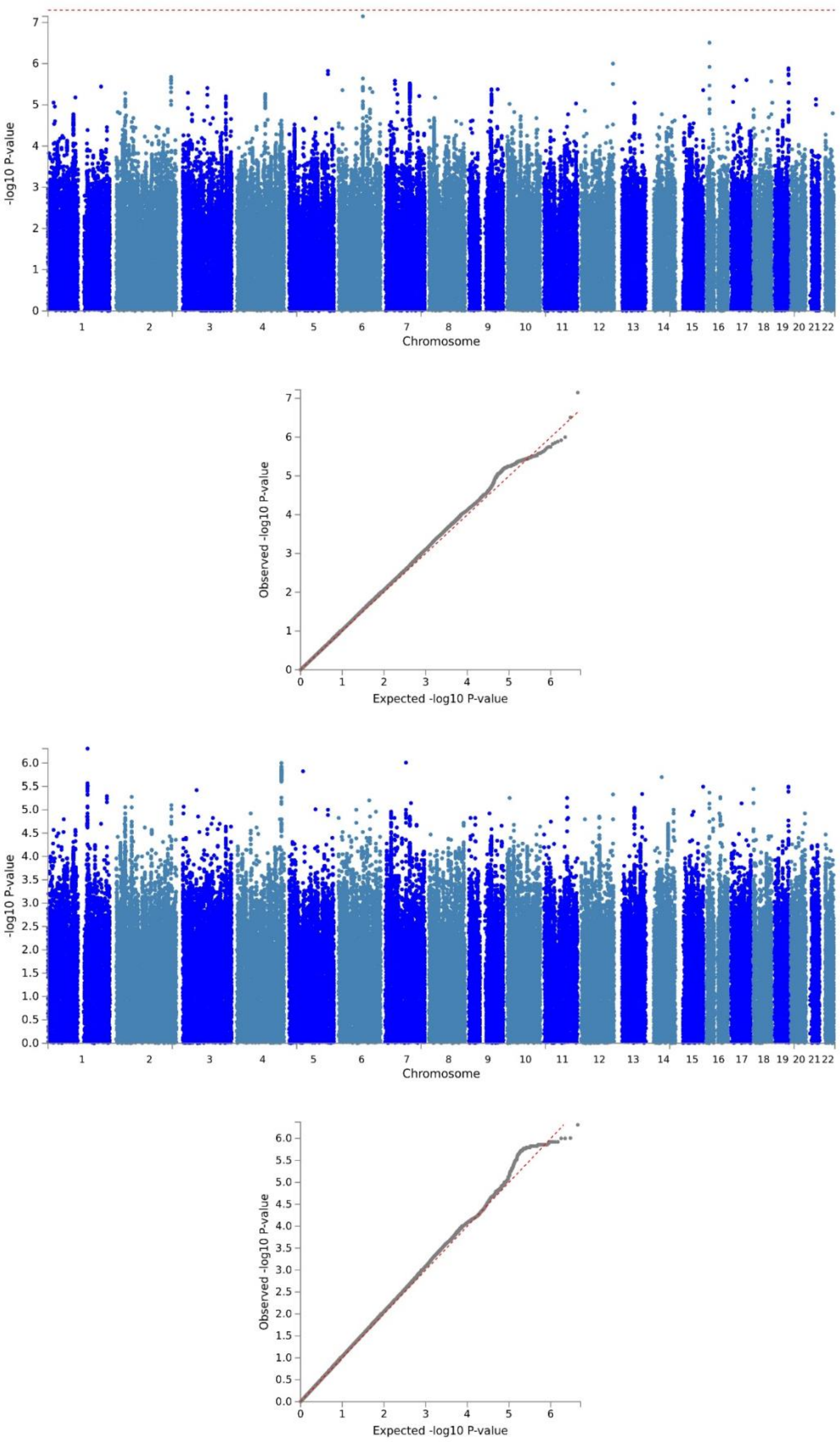

*Supplementary Figure 9.* **Suicidal ideation** Manhattan and QQ plots (binary phenotype above and ordindal below)

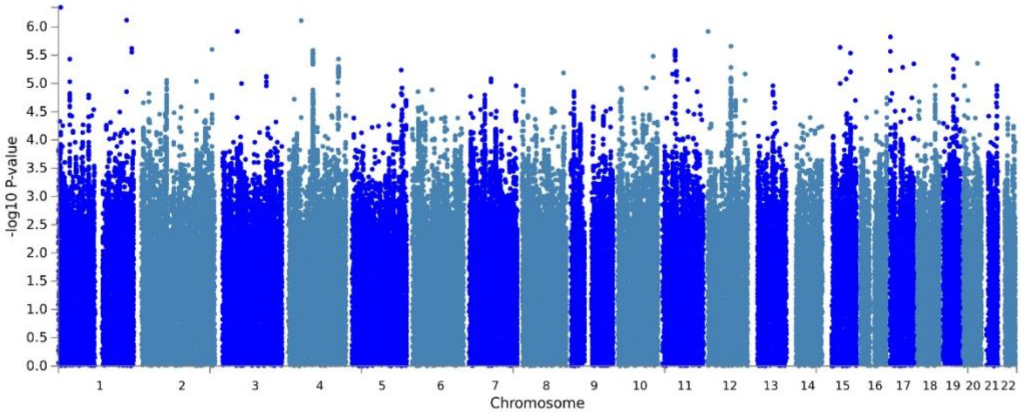


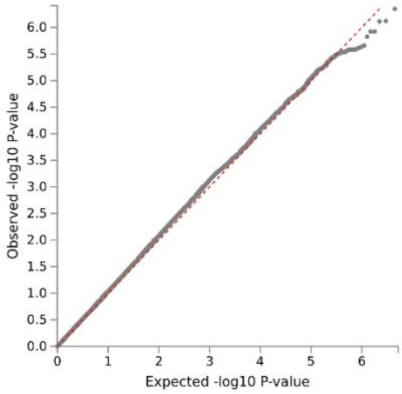


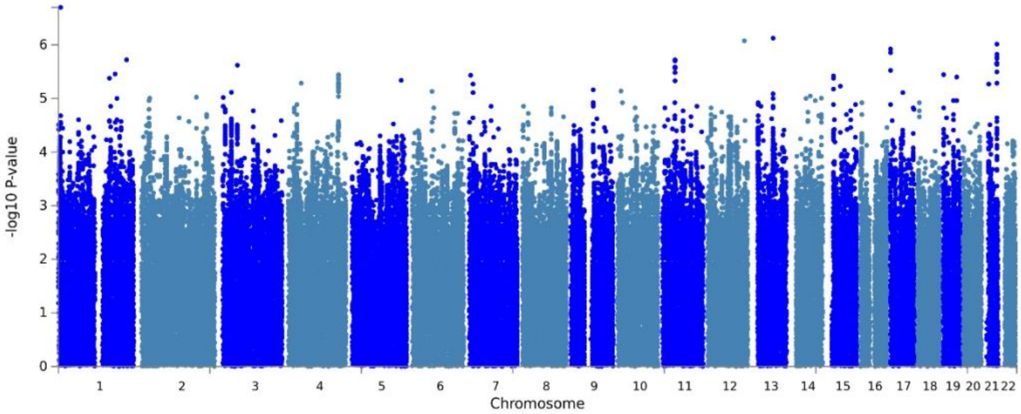


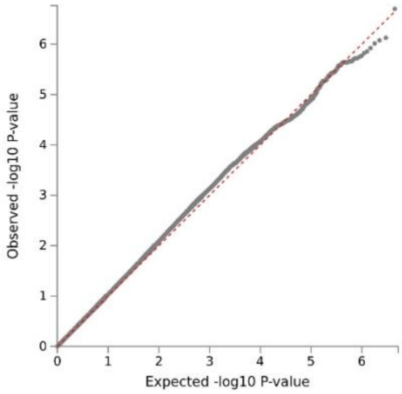

*Supplementary Figure 10.* **Sum-score** Manhsattan and QQ plots (binary phenotype above and ordindal below)

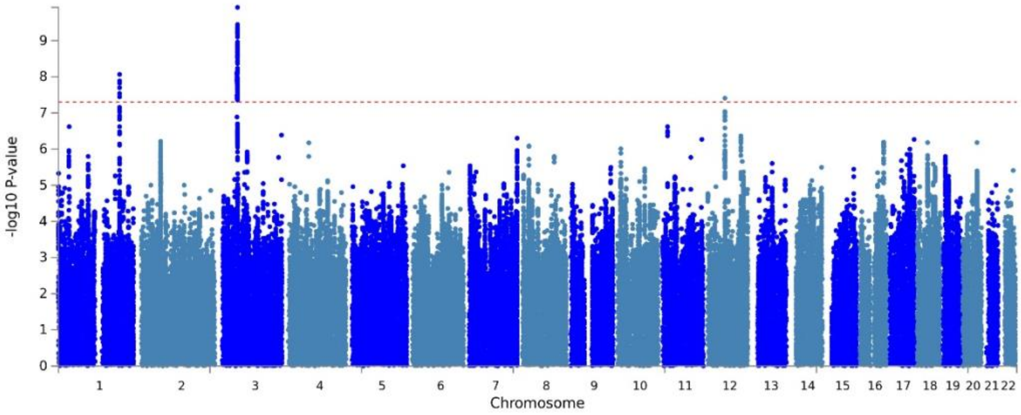


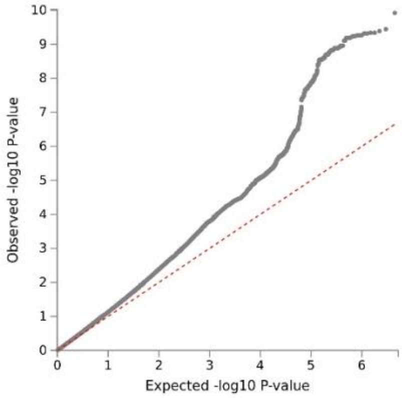


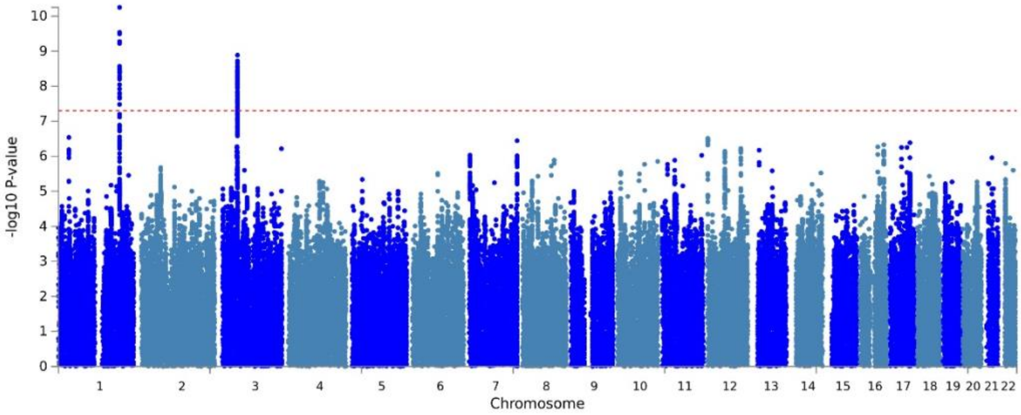


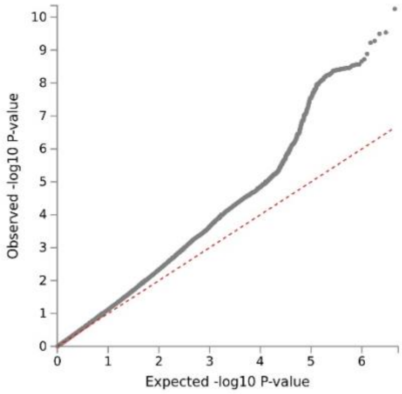

*Supplementary Figure 11.* Comparison of inter-item genetic correlations (rg) between binary items (above diagonal in correlation heatmap) and ordinal items (below diagonal).

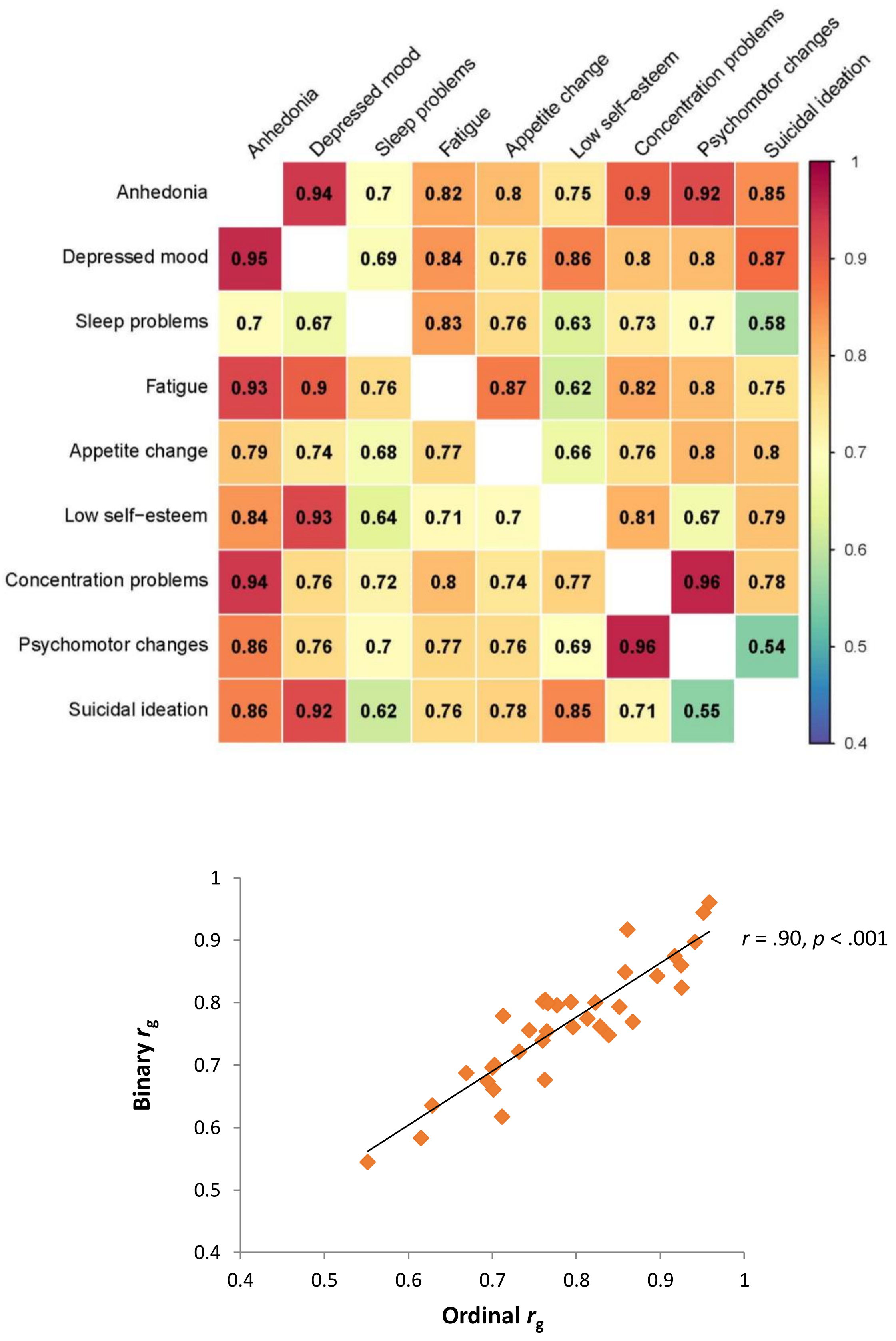

*Supplementary Figure 12.* Scatterplots of genetic correlations ($r_g$) vs. phenotypic correlations ($r_s$) for both ordinal and binary items.

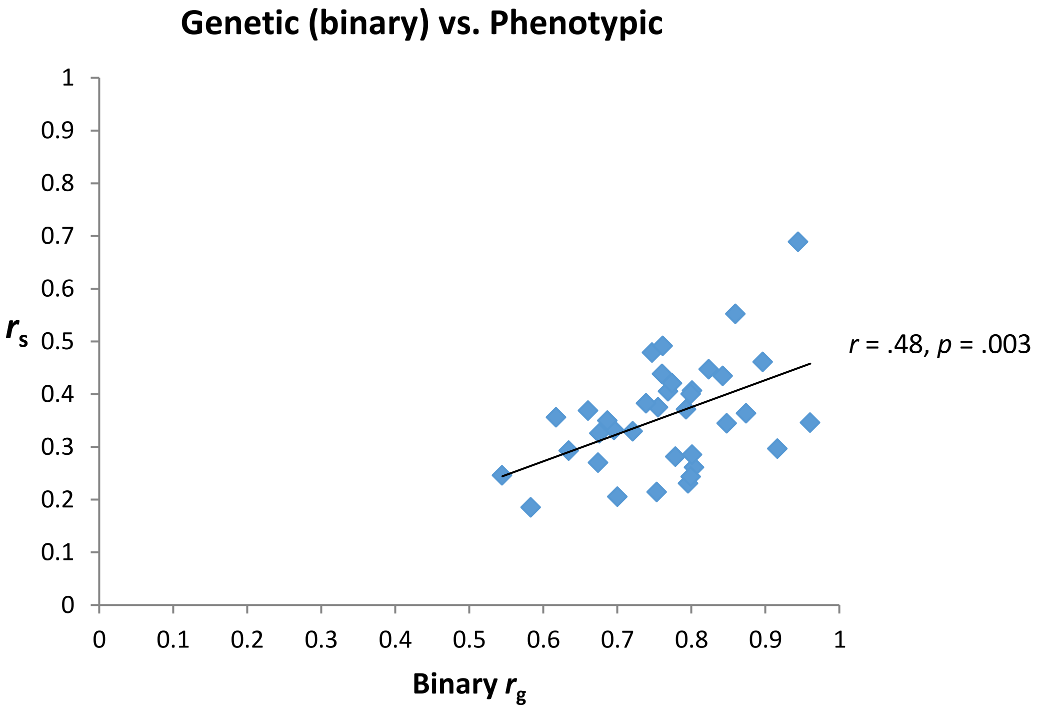


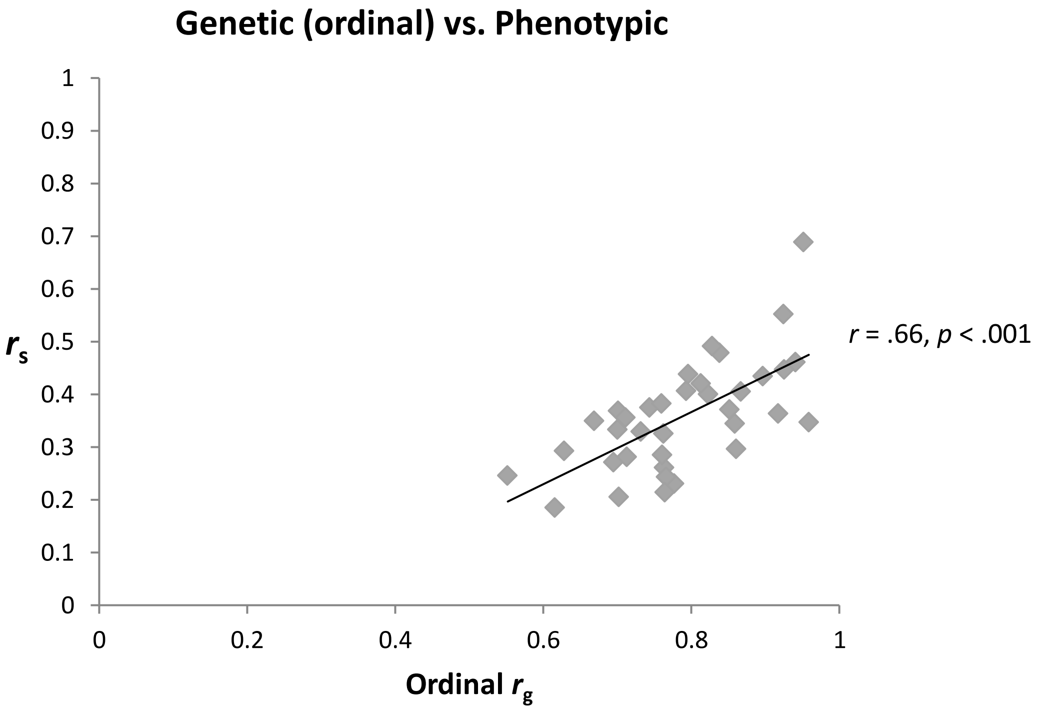

*Supplementary Figure 13.* Flow diagram of exclusions / inclusions leading to final sample size.

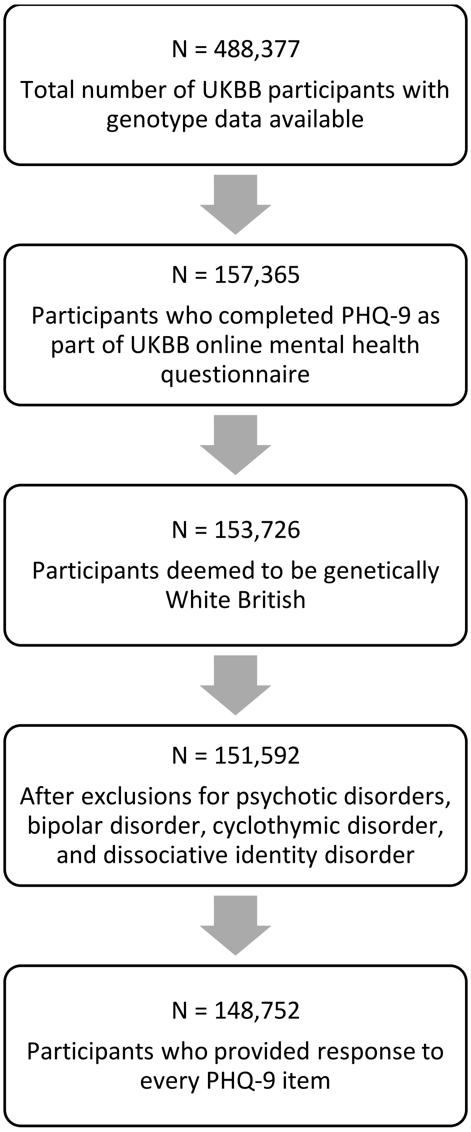
