## Supplementary Methods for "Investigating Genetic Heterogeneity in Major Depression Through Item-level Genetic Analyses of the PHQ-9"

**LDSC analyses**

Single-trait LD Score Regression was used to calculate SNP-based heritability of each of the phenotypes. LD score regression analyses were based on summary statistics obtained from the GWA analyses and pre-computed LD scores of individuals of European ancestry based on 1000 Genomes Project reference data. LD Score Regression is based on the premise that the effect size of a SNP includes the effects of all SNPs in LD with that SNP (Bulik-Sullivan, Loh, et al., 2015; Yang et al., 2011). Therefore, a SNP with a high LD score (a SNP that tags many other SNPs) will on average have a larger effect size than a SNP with a low LD score. By regressing the effect size from the GWA analysis onto the LD Score for each SNP, the slope of the regression line estimates the *h*^2^ _SNP_ of that phenotype. Cross-trait LD Score Regression (Bulik-Sullivan, Finucane, et al., 2015) was used to calculate genetic correlations. The product of effect sizes (z scores) of two traits are regressed on LD Score for each SNP. The genetic covariance between the two traits is estimated from the slope of the regression line and is not biased by sample overlap.

### Missing Data

The amount of missing data for each item of the PHQ-9 is displayed in Table 4. All items had very low proportions of missing data (ranging from 0.14% to 0.76%) and hence were adequately assessed. Suicidal ideation had the highest rate of missing data. A significant Little’s MCAR test suggested that data were not missing completely at random, χ^2^ (1347, *N* = 151,592) = 7209.45, *p* < .001, suggesting a systematic pattern to missing data. Separate variance t-tests were conducted to examine whether the occurrence of missing data could be predicted. Across these tests older individuals were significantly more likely to have a missing response in all items except suicidal ideation.

Frequency of Missing Data for All Items

| Item | N present | N missing | % missing |
| --- | --- | --- | --- |
| Anhedonia | 151,177 | 415 | 0.27 |
| Depressed mood | 151,050 | 542 | 0.36 |
| Sleep problems | 151,266 | 326 | 0.22 |
| Fatigue | 151,264 | 328 | 0.22 |
| Appetite changes | 151,375 | 217 | 0.14 |
| Low self-esteem | 150,853 | 739 | 0.49 |
| Concentration problems | 151,369 | 223 | 0.15 |
| Psychomotor changes | 151,328 | 264 | 0.17 |
| Suicidal ideation | 150,439 | 1,153 | 0.76 |

These findings need be considered under the following context: (1) the present study has a very large sample size, to which both *χ*^2^ and t-tests are sensitive and can inflate type-I error rates (Martin-Löf, 1974); (2) although missing data could be predicted by age, differences in age were small ranging from 1.87 to 4.62 years; and (3) all items had very low rates of missing data (less than 1%). As such, it is unlikely that the presence (or subsequent deletion) of missing data biases the sample. Having the same sample size across all items was preferred in order for additional sum-score phenotypes be evenly created. Therefore list-wise deletion of participants missing any data was done, removing 2,840 individuals from the sample. This represented a less than 2% reduction in sample size having a minimal impact on statistical power.
